# Design of a Novel Multiplex Real Time RT-PCR Assay for SARS-CoV-2 Detection

**DOI:** 10.1101/2020.06.04.135608

**Authors:** Wei Zhen, Gregory J. Berry

## Abstract

Coronavirus Disease 2019 (COVID-19) caused by severe acute respiratory syndrome coronavirus 2 (SARS-CoV-2) has resulted in more than 386,000 deaths globally as of June 4, 2020. In this study, we developed a novel multiplex real time reverse transcription (RT)-PCR test for detection of SARS-CoV-2, with primers designed to amplify a 108 bp target on the spike surface glycoprotein (S gene) of SARS-CoV-2 and a hydrolysis Taqman probe designed to specifically detect SARS-CoV-2. Following our design, we evaluated the Limit of detection (LOD) and clinical performance of this laboratory-developed test (LDT). A LOD study with inactivated whole virus exhibited equal performance to that seen in the modified CDC assay with a final LOD of 1,301 ± 13 genome equivalents/ml for our assay vs 1,249 ± 14 genome equivalents/ml for the modified CDC assay. In addition, a clinical evaluation with 270 nasopharyngeal (NP) swab specimens exhibited 98.5% positive percent agreement and 99.3% negative percent agreement with the modified CDC assay. The multiplex design of this assay allows the testing of 91 patients per plate, versus a maximum of 29 patients per plate on the modified CDC assay, providing the benefit of testing significantly more patients per run and saving reagents during a time when both of these parameters have been critical. Our results demonstrate that our multiplex assay performs as well as the modified CDC assay, but is more efficient and cost effective and is therefore adequate for use as a diagnostic assay and for epidemiological surveillance and clinical management of SARS-CoV-2.

## Introduction

Severe acute respiratory syndrome coronavirus 2 (SARS-CoV-2) was first discovered as etiologic agent of Coronavirus Disease 2019 (COVID-19) in the city of Wuhan, Hubei providence, China by the end of December of 2020 (1), and is the seventh coronavirus known to infect humans and also to be transmitted from human to human. The four seasonal coronaviruses (HKU1, NL63, OC43 and 229E) are associated with mild symptoms, whereas SARS-CoV, MERS-CoV and SARS-CoV-2 can cause severe acute respiratory disease (2,3). SARS-CoV-2 belongs to the betacoronavirus genus and is an enveloped, single-strand RNA virus with a ^~^29.8 kb genome, which can cause a wild range of clinical presentations from asymptomatic or mild illness to fatal outcomes (4,5). Furthermore, the symptoms of COVID-19 patients can be similar to that of patients with other seasonal respiratory infections. Presently, there are no available specific therapeutics or vaccinations against COVID-19, making early and accurate diagnosis for this very contagious disease the key mitigation strategy.

Nucleic Acid Amplification Test (NAAT)-based assays for detection of SARS-CoV-2 in respiratory specimens have been the standard diagnostic method. To date, the FDA has issued 34 lab developed COVID-19 molecular assays <u>E</u>mergency <u>U</u>se <u>A</u>uthorization (EUA) (6). The US CDC has developed the most widely used SARS–CoV-2 assay which includes primers and probes to detect the N1 and N2 regions of the nucleocapsid gene and also the human RNase *P* gene to monitor RNA extraction and ensure specimen quality (7). For each specimen performed on the modified CDC assay, there are three Master Mix sets, including N1, N2 and RNase P. Each master mix needs to be independently prepared and dispensed into the appropriate wells before the extracted clinical sample RNA is added to each well. The modified CDC assay has been shown to have high analytical sensitivity and ideal clinical performance when compared to three commercially available COVID-19 diagnostic platforms issued EUA status by the FDA (8).

The aim of our current study was to develop and then evaluate the analytical sensitivity and clinical performance of an efficient and cost effective laboratory-developed test (LDT) on the 7500 Fast Dx real time PCR instrument. To that end, we developed an LDT that targets the S gene of SARS-CoV-2 and compared it’s clinical performance to the modified CDC assay for the detection of SARS-CoV-2 in nasopharyngeal (NP) specimens from individuals suspected of potentially having COVID-19.

## Materials and Method

### Primers and probe design

Available whole genome sequence of SARS-CoV-2 (as of February 27, 2020) retrieved from the NCBI GenBank database and the Global Initiative on Sharing All Influenza Database (GISAID) were aligned using Clustal Omega software from EMBL-EBI. The primers and probe were designed using Primer Express 3.0 software in the S gene of SARS-CoV-2 and were synthesized by Integrated DNA Technologies, Inc. (IDT). In addition, the primers and probe of human RNase P gene used for the assay internal control were also synthesized by IDT and were the same sequences used in the CDC assay for this gene target (7). Here, the 5’ base of probe of RNase P gene was modified and labeled with Cy5, and the 3’ base of probe was labeled with Black Hole Quencher 2(BHQ2) to allow multiplexing of our LDT assay.

### Study design

A modified version (v3) of the CDC assay was used as a reference method in this study (8). A total of 270 NP specimens (130 positive and 140 negative specimens) originally submitted for SARS-CoV-2 testing at Northwell Health Laboratories between March and April 2020 were selected for this study. The 270 specimens were initially tested by the modified CDC assay and extracted RNA was stored at −80°C until testing with our LDT was performed. The specimens were selected as any consecutive specimen that was performed on the modified CDC assay during the timeframe of this study and represented our true positivity rate (^~^50%), including positive specimens spanning the range of positivity levels, including 24 specimens with low viral load that characterized by high Cycle threshold (Ct) value in range of 31.7-39.7 by the modified CDC assay. For discordant results, molecular testing was repeated from RNA extraction on both assays. This study was performed in order to validate this LDT for clinical use.

### RNA extraction

Total RNA was extracted from 110 μl of patient NP specimen by the NucliSENS easyMag platform (BioMérieux, Durham, NC) according to the manufacturer’s instructions, and the final elution volume was 110 μl. In order to monitor the extraction process, a Negative Extraction Control was included in each extraction run.

### Laboratory developed real time RT-PCR assay specific for SARS-CoV-2 detection

In this one-step, qualitative real time RT-PCR assay, TaqPath 1-step RT-qPCR kit (Catalog no. A15299, Thermo Fisher Scientific) was used to perform cDNA synthesis and PCR amplification on the 96-well plate at a 20 μl final reaction volume. The per reaction mix contained 5 μl of 4X RT-PCR Master Mix, 0.72 μl of S gene forward primer at 25 μM, 0.72μl of S gene reverse primer at 25 μM, 0.16 μl of S gene probe at 25μM, 0.64 μl of RNase P gene forward primer at 25 μM, 0.64 μl of RNase P gene reverse primer at 25 μM, 0.16 μl of RNase P probe at 25 μM, 6.96 μl of nuclease free water and 5 μl of extracted RNA. The thermal cycler profile consisted of 25°C for 2 min, 50°C for 15 min, 95°C for 2 min and followed by 45 cycles at 95°C 3 sec, 55°C for 30 sec, and was conducted on the 7500 Fast Dx real time PCR instrument. Each run included a No Template Control, Negative Extraction Control and a SARS-CoV-2 positive control. The LDT used the same RT-PCR reagents and conditions as the modified CDC assay, except for the target primer and probe.

### Analytical Sensitivity (Limit of Detection)

Limit of detection (LOD) was determined by extracting and testing serial dilution panels of quantified inactivated SARS-CoV-2 from Isolate USA-WA1/2020 (NR-52287, BEI Resources, Manassas, VA) SARS-CoV-2 viral material was provided at a concentration of 4.1 x 10^9 genome equivalents (GE)/ml, from which the following serial dilutions were prepared in GE/ml: 4000, 2000, 1000, 500, 250, 125 using Ambion^®^ RNA Storage Solution (Catalog No. AM7001, ThermoFisher Scientific) to prevent the potential RNA degradation, and replicates ranging from 4-10 at each dilution went through nucleic acid extraction on different days and were tested on both the modified CDC assay and our LDT. LOD was defined as the concentration of the lowest dilution that can be detected with greater than 95% probability and was determined by Probit analysis.

### Analytical Specificity

The specificity of our LDT primers and probe for SARS-CoV-2 detection was evaluated by *in silico* analysis and by testing a SARS-CoV control (GenBanK: MG772933.1) (Cat no. 10006624, IDT), MERS-CoV control (GenBank: MK796425.1)(Cat no. 10006623, IDT) and a panel of respiratory pathogens covering Coronavirus (229E, NL63, OC43, HKU1), Influenza A, H3, Influenza B, 2009 H1N1, RSV A, RSV B, Parainfluenza virus type 1-4, Human metapneumovirus, Adenovirus, *Mycoplasma pneumoniae*, and *Chlamydia pneumoniae* that were initially identified using a multiplex respiratory panel at Northwell Health laboratories.

### Statistical methods

The final result interpretation algorithm for reporting a positive specimen requires both N1 and N2 targets to be detected in the modified CDC assay (7), and both of the modified CDC assay and our LDT use a Ct < 40 as the criterion for positivity. Percent positive agreement (PPA), percent negative agreement (NPA), Kappa, and two-sided (upper/lower) 95% confidence interval (CI) were calculated using Microsoft^®^ Office Excel 365 MSO software (Microsoft, Redmond, WA). As a measure of overall agreement, Cohen’s kappa values (κ) were calculated, with values categorized as follows: >0.90 = almost perfect, 0.90 to 0.80 = strong, 0.79 to 0.60 = moderate, 0.59 to 0.40 = weak, 0.39 to 0.21 = minimal, 0.20 to 0 = none (9, 10).

## Results

### Design of primers and probe for SARS-CoV-2 detection

Using primer and probe design tool, the assay primers and probe specifically targeting S gene of SARS-CoV-2 (Table 1) were designed to amplify a 108 bp target on the conserved S gene based on multiple sequence alignment and *In silico* analysis for our LDT primers and probe was performed. The primers and probe designed for S gene detection were designed using all available SARS-CoV-2 whole genome sequences. Our analysis of sequence alignments revealed that region of the S gene of SARS-CoV-2 targeted by designed primer and probe set have 100% similarity with all available SARS-CoV-2 whole genome sequence from GenBank and GISAID database at the time of development.

**Table 1:**
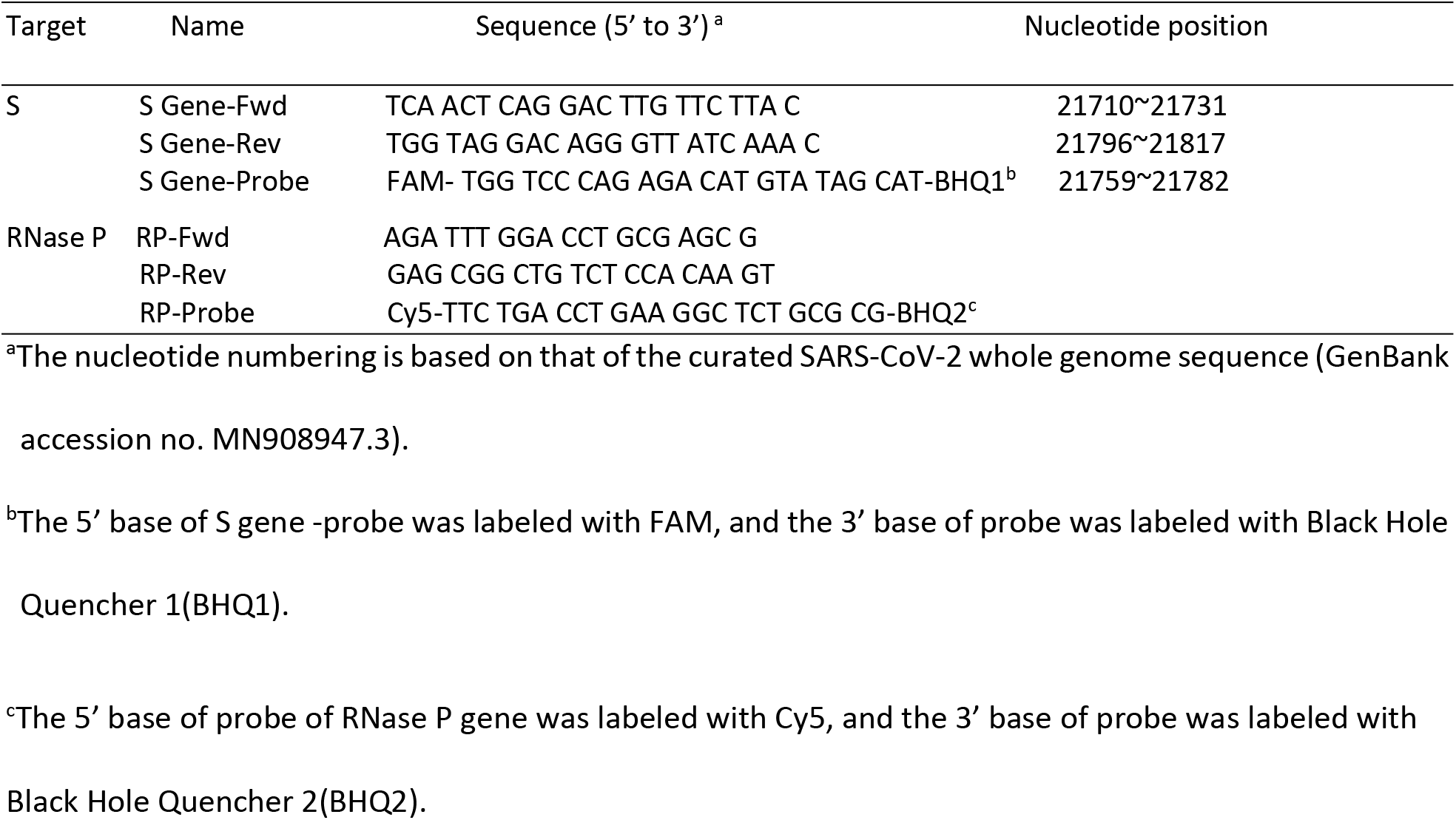
Real time RT-PCR primer / probe set of S gene target for specific detection of SARS-CoV-2 and primer / probe set of RNase P gene.

### Analytical Sensitivity

Quantified inactivated SARS-CoV-2 virus particles were used to prepare serial dilutions (4,000 GE/mL to 125 genome equivalents (GE)/ml, in 2-fold dilutions) to determine the LOD of our LDT as well as the modified CDC assay. The LOD was defined as the lowest dilution in which all replicates were detected with 95% positivity rate by Probit analysis. The LOD of LDT was 1301±13 GE/ml for the S gene target. For the modified CDC assay, the LOD was 1249±14 GE/mL for the N1 target and 946±11 GE/ml for the N2 target (Table 2). The final LOD of modified CDC assay was 1249±14 GE/mL in accordance with the result interpretation algorithm.

**Table 2:** Summary of limit of detection results

| Molecular assay | target | No. of replicates detected / total no. of replicates at each dilution expressed by GE /ml<sup>a</sup> (% positive rate) |  |  |  |  |  | Probit (±95% CI)<sup>b</sup> GE/ml | Final LOD<sup>c</sup> GE/ml |
| --- | --- | --- | --- | --- | --- | --- | --- | --- | --- |
|  |  | 4000 | 2000 | 1000 | 500 | 250 | 125 |  |  |
| LDT<sup>d</sup> | S | 4/4 | **6/6**<br>**(100)** | 9/10<br>(90) | 5/10<br>(50) | 2/8<br>(25) | 1/8<br>(25) | 1301±13 | 1301±13 |
| Modified CDC assay | N1 | 4/4 | **6/6**<br>**(100)** | 9/10<br>(90) | 7/10<br>(70) | 6/8<br>(75) | 1/8<br>(25) | 1249±14 | 1249±14 |
|  | N2 | 4/4 | 6/6<br>(100) | **10/10**<br>**(100)** | 5/10<br>(50) | 2/8<br>(25) | 1/8<br>(25) | 946±11 |  |

<sup>a</sup>GE/ml: genome equivalent/ml.
<sup>b</sup>95% CI, upper/lower (±) 95% confidence interval.
<sup>c</sup>The final LOD were based on the both assay's results interpretation algorithm and determined by the
Probit analysis
<sup>d</sup>LDT: Laboratory developed test.

### Analytical specificity

The analytical specificity of our LDT was evaluated using *in silico* analysis and also by testing a panel of common respiratory viruses, including the four strains of circulating coronaviruses. *In silico* analysis and Blastn analysis were performed against the standard and betacoronavirus database of National Center for Biotechnology Information (NCBI) and results showed no cross reactivity with other respiratory pathogens; A panel of respiratory pathogens was also tested to further evaluate the specificity of our LDT. According to our results (Data not shown), a specificity of 100% was achieved and our LDT had no cross-reaction with any of the pathogens tested.

### Clinical performance

The clinical performance of our LDT was evaluated by comparing to the modified CDC assay. A total of 270 clinical NP specimens from standard of care testing for SARS-CoV-2 on the modified CDC assay from March 2020 to April 2020 at Northwell Health Laboratories were tested. The LDT demonstrated a PPA of 98.5% (95% CI, 0.946-0.996), and NPA of 99.3% (95% CI, 0.961-0.999), with the overall percent agreement between both assays being 98.9%. A kappa value of 0.978 (95% CI 0.953-1.0) indicated perfect agreement (Table 3). For discordant results from three specimens, two specimens were detected by the modified CDC assay and were not detected by LDT and had a Ct value of 37.3 for N1, 39.6 for N2 and 38.2 for N1 and 39.2 for N2, respectively. Following a fresh RNA extraction and retesting, the specimen with 37.3 N1 and 39.6 N2 Ct values was detected by both assays, and the other specimen resulted as not detected by both assays. One specimen was initially detected by LDT and not by the modified CDC assay, exhibiting a Ct value of 32.0 on the LDT. After retesting, both the modified CDC assay and the LDT assay were positive.

**Table 3:**
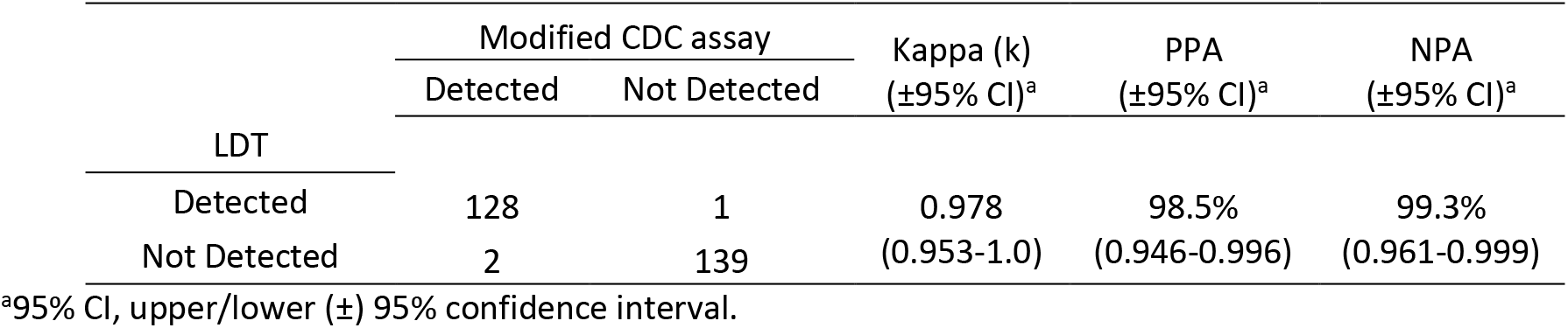
Clinical performance comparison of between our LDT and modified CDC assay for the detection of SARS-CoV-2 RNA (n=270)

## Discussion

In the present study, a multiplex real time RT-PCR assay was developed and validated for SARS-CoV-2 specific detection in NP specimens on the 7500 Fast Dx real time PCR instrument.

The findings demonstrate that our LDT design has comparable clinical performance for the specific detection of SARS-CoV-2 RNA in NP specimens and is more efficient and cost effective in comparison to the modified CDC assay. Our LDT also showed significant advantages over the modified CDC assay, since only one set of primer and probe Master Mix is required to prepare and dispense per specimen, in contrast to three sets of Master Mix preparation and the use of three wells for each patient specimen by the modified CDC assay. Our multiplex design allows us to run 91 patients per plate, versus a maximum of 29 patients per plate on the modified CDC assay. Overall, this allows us to run more than 3 times as many patients per run and also adds to the ease of setting up each run. Additionally, the saving of reagents and consumables is another advantage at a stage where there are currently global shortages of reagents and major assay supply chain issues.

The design of the primers and probe for our LDT is based on multiple sequence alignment of all SARS-CoV-2 genome sequences that were available between January 11 and February 27 of 2020. Since RNA viruses are well known for its high mutation and recombination rates (11), we wanted to confirm that there were no significant new mutations of the region of S gene of SARS-CoV-2 that is the target in our assay that could potentially affect assay performance. To this end, we analyzed an additional 140 SARS-CoV-2 genome sequences uploaded after February 27, 2020 to GenBank and the GISAID database from different countries and performed an alignment with Clustal Omega. This alignment showed that the forward primer and probe are conserved (with 100% homology) to the S gene target regions of the SARS-CoV-2 sequences. One exception was seen with the reverse primer, a single base mismatch of S gene target region in one sequence (MT385417.1) from a total of 240 SARS-CoV-2 sequence analyzed both before and after late Febuary. This one mismatch is questionable, since it is not in keeping with the other sequences available and the databases are not curated, and therefore occasionally contain errors.

Limitations of our study include that our LDT is a single site evaluation at Northwell Health Laboratories. In addition, we only use a single target gene for SARS-CoV-2 detection. While there has been a trend toward dual-target design in commercial assay for the detection of pathogens (8,12), occasional monitoring of SARS-CoV-2 sequences to verify that mutations have not developed in the region targeted by our primers and probe is an adequate quality monitor to ensure continued consistent analytical performance.

In summary, our LDT has comparable analytical sensitivity and accuracy for specific detection of SARS-CoV-2 RNA when compared to the modified CDC assay. In addition, it also showed superior efficiency and cost-effectiveness and can be somewhat easily-established in other laboratories. These findings make our novel multiplex SARS-CoV-2 assay a suitable alternative for the accurate diagnosis of SARS-CoV-2, with the added benefit of superior efficiency and cost-effectiveness.

